# A Novel Optical Design Enabling Lightweight and Large Field-of-View Head-Mounted Microscopes

**DOI:** 10.1101/2021.09.03.458947

**Authors:** J. R. Scherrer, Galen F. Lynch, Jie J. Zhang, Michale S. Fee

**Author notes:** equal contribution (J. R. Scherrer, Galen F. Lynch).

## Abstract

We present a novel fluorescence microscope light path that enables imaging, during free behavior, of thousands of neurons in mice and hundreds of neurons in juvenile songbirds. The light path eliminates traditional illumination optics, allowing for head-mounted microscopes that have both a lower weight and a larger field-of-view (FOV) than previously possible. Using this light path, we designed two microscopes: one optimized for field-of-view (∼4 mm FOV; 1.4 g), and the other optimized for weight (1.0 mm FOV; 1.0 g).

## Main Text

Miniature head-mounted microscopes have expanded the neuroscience toolkit by enabling *in vivo* optical measurements of neural activity in freely moving animals^1-3^. Such recordings have been used to study the neural basis of behaviors that are difficult or impossible to recapitulate in head-fixed preparations^4^ like social interaction^5^, vocalization^6^, sleep^7^, and navigation^8^. Despite the impact that head-mounted fluorescent microscopes have had in neuroscience, challenges remain in extending this approach to large populations of neurons and to small model organisms. Understanding how behavior is controlled by neural populations across brain regions requires simultaneous recording in thousands of neurons, which until now has only been possible in small animals using head-fixed preparations^9,10^. Performing these experiments in freely moving animals requires microscopes with a larger field-of-view (FOV) than existing technologies, which are limited to recording from several hundred individually resolvable neurons due to their limited FOV. At the same time, imaging in small organisms such as juvenile mice and juvenile songbirds requires devices with lower weights than existing microscopes, which are too heavy to be used in these small animals. Radically new approaches such as computationally reconstructed imaging are a promising potential solution to FOV and weight limitations^11,12^, but have yet to be successfully applied to functional brain imaging. More incremental improvements to microscope designs have both decreased device weight and increased FOV^13-17^, but these devices are still limited by design tradeoffs that constrain either weight or optical performance.

To address these challenges and opportunities, we describe a novel optical pathway that allows for much lighter microscopes with larger fields of view than previously possible, reducing the tradeoff between weight and FOV inherent to previous designs. Most epifluorescence microscopes couple excitation light using a dichroic filter placed between the objective lens and tube lens (Fig. 1a). The added path length due to the dichroic not only increases microscope dimensions, but also necessitates a larger tube lens to accommodate off-axis rays. The large tube lens size means that the complex optics required for cellular resolution and large FOV imaging quickly become prohibitively heavy. Alternatively, microscopes can be optimized for low device weight, but they typically have a limited FOV (around 1 mm^2^), lower resolution, or decreased light-gathering ability. In contrast, our new optical path brings the objective and tube lens closer together, allowing the optical assembly to be more compact without sacrificing optical performance. To do so, we eliminate the dichroic and instead introduce excitation light at the back aperture of the objective lens (Fig. 1b) with a low-profile light guide and coupling prism.

**Figure 1.**
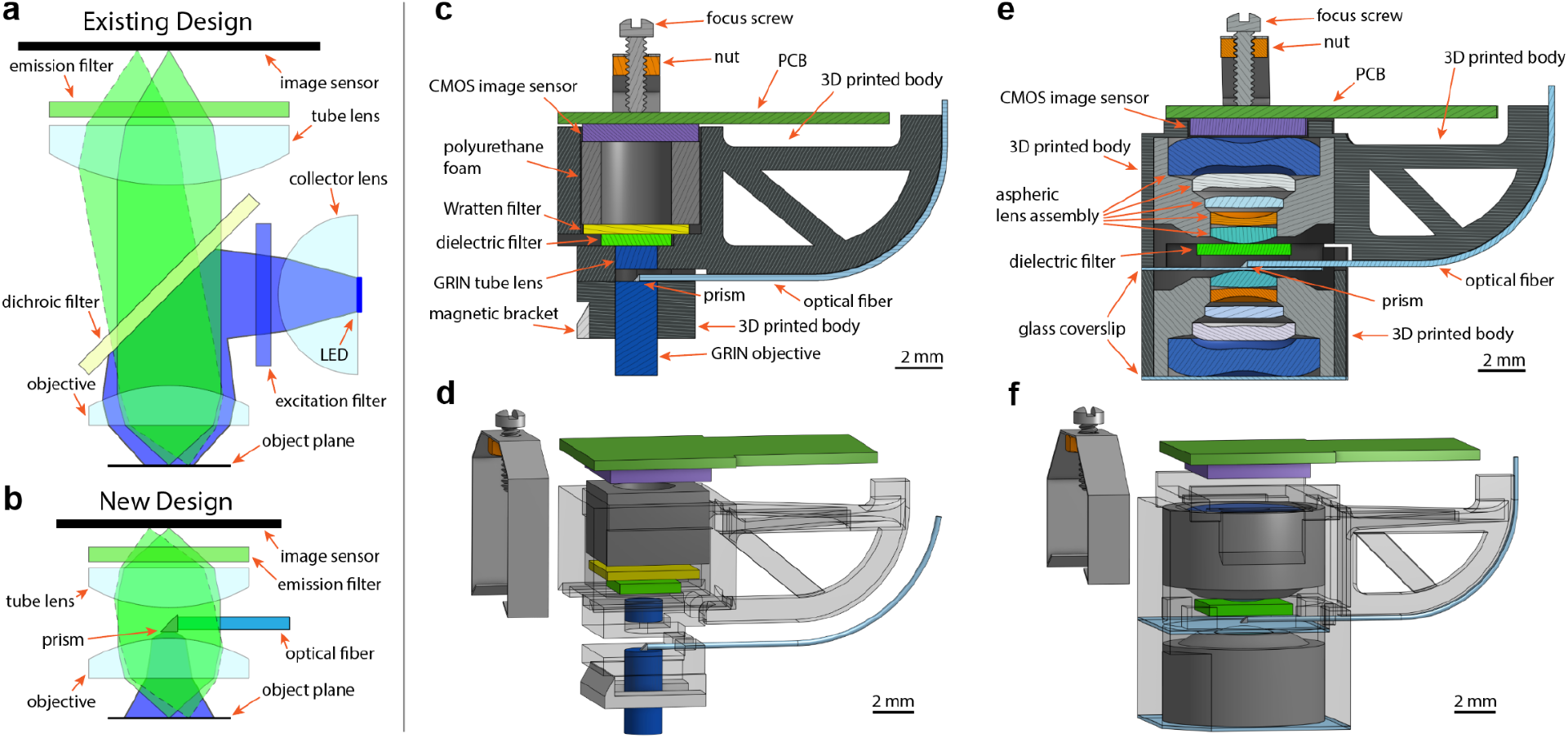
Design of head-mounted miniature microscopes. **a**, Schematic representation of optical path for standard head-mounted microscopes and **b**, for the microscopes in this paper. Ray traces are the excitation light path (blue) and emission light paths for points on-axis (solid green) and off-axis (dashed green). **c**, cross-section and **d**, explosion diagrams for the Featherscope. **e**, cross-section and **f**, explosion diagrams for the Kiloscope.

We used our new light path to design two head-mounted microscopes that advance the state-of-the-art in device weight and FOV. The first design, the Featherscope, is optimized for the lowest possible weight without sacrificing resolution. It weighs 1.0 g but does not sacrifice cellular resolution across a 1.0 × 1.0 mm FOV, enabling neural recordings in animals that are too small for existing head-mounted microscopes. We used this first microscope to report, for the first time, functional calcium imaging in freely behaving juvenile songbirds during song learning. The second design, the Kiloscope, is optimized for maximum FOV while still achieving an extremely low weight. It provides cellular resolution imaging across a 4.8 × 3.6 mm FOV in a device weighing only 1.4 g, allowing us to simultaneously record from thousands of neurons in a freely behaving mouse.

The Featherscope provides cellular resolution imaging in such a lightweight and compact package by combining the novel optical path described above with compact gradient index (GRIN) optics for both the objective and tube lenses (Fig. 1c,d), a lightweight focusing mechanism, and a custom high-density image sensor printed circuit board (PCB). GRIN lenses both reduce device weight and outperform single spherical lens elements (Supp. Fig. 1a). Excitation light is separated from emission light using a combination of an interference filter and an angle-independent organic dye Wratten filter. Instead of using bulky focusing mechanisms such as electrowetting lenses or screw-tightened sleeves, we provide focus adjustment by compressing the image sensor PCB against polyurethane foam using a screw bracket, allowing fine adjustment of the focus over a range of approximately 200 μm.

The Kiloscope achieves a wide field of view by using the same optical pathway in combination with precision molded plastic aspheric lens assemblies developed for smartphone cameras (Fig. 1e,f). These lens assemblies achieve diffraction-limited resolution across several millimeters in a compact package. Using two such assemblies for both the objective and tube lenses in a “double macro” configuration^18^, we achieve high resolution imaging with near-flat field curvature over a large FOV (Supp. Fig. 1b). The high chief ray angle of the lenses was compensated for using an image sensor with a shifted microlens array.

We provide illumination for both microscope designs using a fiber-coupled 473 nm laser, which allows for a higher power density than an onboard LED. To reduce imaging artifacts caused by laser speckle, we modulate the laser output using an oscillating ground glass diffuser (Supp. Fig. 2). The despeckled laser light is then coupled into a flexible light guide that terminates at a 300 μm 90**°** prism mounted to the back aperture of the objective lens. The prism-fiber assembly partially occludes the optical path, reducing the overall light collection efficiency by approximately 20%. Such back-aperture occlusion has a minor effect on diffraction limited spot size, and produces a slight increase in vignetting, but preserves the fine detail necessary for neural imaging. We compensate for optical vignetting (Supp. Fig. 3) by increasing the laser illumination intensity at the edges of the FOV relative to the center using a pair of conical lenses placed before the fiber input coupling optics (Supp. Fig. 2).

Both microscope designs use a custom printed circuit board that supports a CMOS image sensor, serializer, and microcontroller. The microcontroller sequences the start-up of the image sensor, acts as a communications bridge between the image sensor and serializer, and digitizes up to two analog inputs (such as electrophysiological recordings) at 10 bits of resolution with a combined sampling rate of 51 kS/s which is transmitted in parallel with video data (Supp. Fig. 6). Video data is transmitted to a computer using an interface board based on the UCLA miniscope design^3^. For long-term recordings in behaving animals, a custom commutator transmits laser illumination light along with video data, electrophysiology data, and power.

We quantified imaging performance using standard resolution test targets and measurements of the point spread function. The Featherscope has a resolution of approximately 4 μm at the center of its field of view, which decreases to 7 μm at the edges (Fig. 2a). The resolution is primarily limited by the astigmatism of the GRIN lenses (Supp. Fig. 1a). The Kiloscope has a resolution of approximately 4 μm at the center of its field of view, which decreases to about 5 μm at the edges (Fig. 2d), and is primarily limited by the pixel size of the image sensor we use. We also used a higher resolution image sensor to test the Kiloscope optics alone and achieved 2 μm resolution (Supp. Fig. 5). Both microscope designs had cellular resolution in neural tissue across the entire FOV (Fig. 2) and sufficiently small field curvature for typical imaging depths (Supp. Fig. 4).

**Figure 2.**
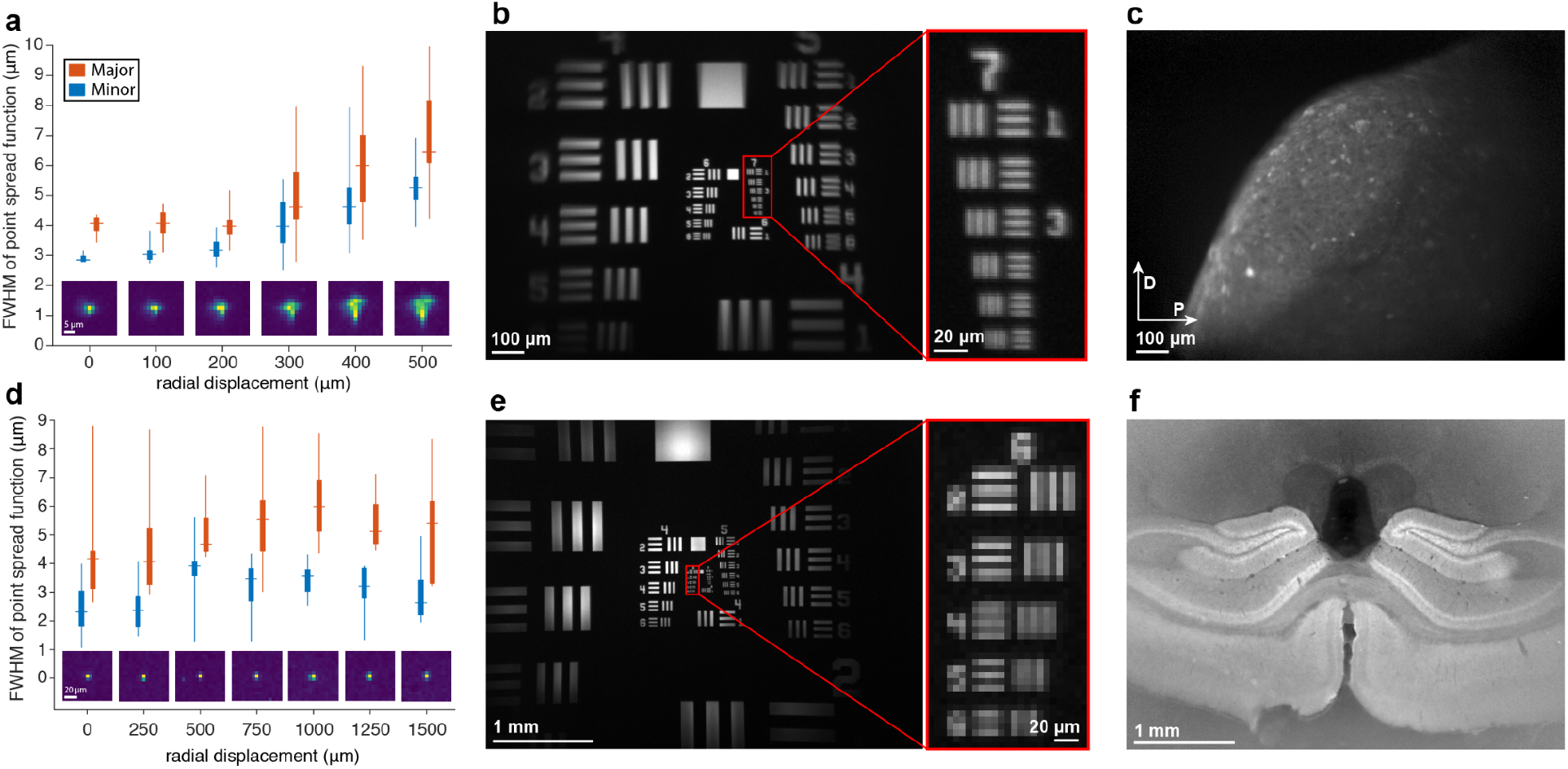
Optical performance measurements. **a**, Measurements of the Featherscope’s point spread function using 1 μm fluorescent beads. Orange and blue data points are measured along the major and minor axes, respectively. The horizontal line represents the median, box edges are the 25th and 75th percentile, and the vertical line extends to the minimum and maximum data points. Inset images are of a representative bead at each position in the object plane. **b**, Image taken with the Featherscope of a USAF 1951 resolution target. The line widths of the largest and smallest elements visible in the inset are 3.91 μm and 2.19 μm, respectively. **c**, Image taken with the Featherscope of zebra finch brain slice, fixed and stained with anti-GFP antibody. The image is corrected for vignetting. **d**, Same as in **(a)** but with the Kiloscope. **e**, Same as in **(b)** but with the Kiloscope. The line widths of the largest and smallest elements visible in the inset are 6.96 μm and 4.38 μm, respectively. **f**, Image taken with the Kiloscope of a fixed coronal brain slice from a Tg(Camk2a-cre)T29-1Stl x Ai95(RCL-GCaMP6f)-D mouse. The image is corrected for vignetting.

To demonstrate the utility of the Featherscope in small animals, we recorded calcium activity in three freely behaving juvenile zebra finches during the subsong and protosyllable stages of song production^19^. Such imaging experiments were previously difficult due to the small size of these young birds, whose singing behavior is significantly impacted by implanted devices weighing more than 1.5 grams. We targeted our recordings to the lateral magnocellular nucleus of the anterior nidopallium (LMAN), a brain region necessary for the earliest stages of song production^19^. After virally expressing GCaMP6f in LMAN, we implanted a GRIN relay lens-prism assembly at the anterior edge of LMAN and let each bird recover. We then installed the Featherscope on the bird’s head and allowed the bird to freely behave for several days while recording from LMAN triggered on song (Supp. Video 1). Implanted birds sang between 3,861-23,545 syllables per day, within the normal range and far exceeding the minimum amount needed for analysis of neural activity^19^. We analyzed the data using EXTRACT, a robust estimation algorithm for extracting microendoscopic calcium activity^20^, and extracted 177, 46, and 225 manually curated footprints that exhibited the spatial profile and temporal fluctuations characteristic of neuronal calcium signals in cortex^23^ (Fig. 3i-l and Supp. Fig. 12). The activity patterns of almost all footprints recapitulate the sparse random bursting seen in previous observations of LMAN activity during song (Fig. 3k,l)^21^.

**Figure 3.**
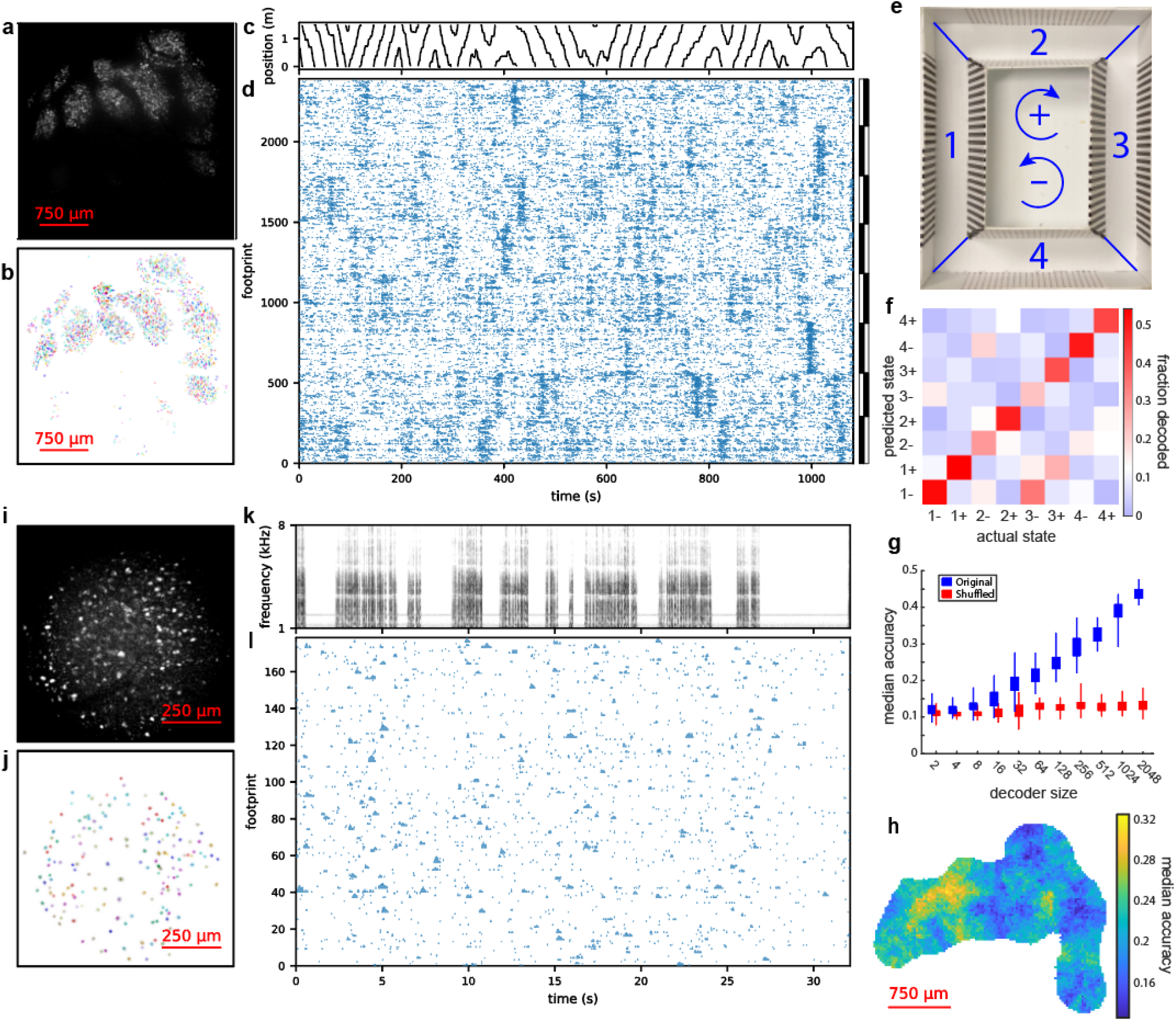
Demonstration of microscopes in awake freely-behaving animals. **a**, Maximum intensity projection of fluorescence signal from AAV1-syn-jGCaMP7f expression in mouse cortex imaged during exploration of a circular maze. The anterior portion of the cranial window is at the top, and the medial portion is to the right. **b**, Spatial extents of 2382 extracted footprints that were identified as putative neurons in the raw video. **c**, Linear position along track and **d** corresponding calcium activity for each of the footprints in **(b)**, sorted by NMF component indicated by vertical bars (see Methods). **e**, Diagram of circular maze used for **(c-d)**, with quadrants and directions labeled. **f**, Decoding matrix for a set of binary SVM models trained to predict the labels in **(e)** based on the neural activity in **(d)**. White is set at chance level. **g**, Decoding accuracies for a series of decoders trained on random subsets of footprints of different sizes, using both the original data (blue) and temporally shuffled activities (red). **h**, Map of decoder accuracies using the 64 closest footprints less than 270 μm to a given point. Colorbar minimum is set at chance level. **i**, Maximum intensity projection of fluorescence signal from AAV9-CAG-GCaMP6f expression in the LMAN of a 49-day old zebra finch imaged during song production. The dorsal portion of the FOV is at the top, and the medial portion is to the left. **j**, Spatial footprints of 177 footprints extracted using EXTRACT. **k**, Spectrogram of a single bout of juvenile song and **l**, corresponding calcium activity for each of the footprints in **(j)** (see Methods).

To demonstrate the performance of the Kiloscope, we used the microscope to record in four freely moving mice and were able to simultaneously record more than a thousand neurons in cortex in multiple mice (Fig. 3a-d and Supp. Fig. 11). For these experiments, we stereotactically located a region of the mouse brain at coordinates associated with primary visual and somatosensory regions, performed approximately twenty viral injections of GCaMP7f across a 4 mm diameter region of cortex, and then implanted a 4 mm diameter glass window in the skull. After letting each mouse recover, we installed the Kiloscope on the head and allowed it to freely explore a circular maze (Fig. 3e) while recording calcium activity for several minutes (Supp. Video 2). We then analyzed the data using EXTRACT and extracted 2948, 401, 1029, and 112 footprints in these mice. In the data from the mouse with the most footprints, manual curation using the same criteria described above yielded 2382 footprints. Lateral shifts of the image due to mouse motion were small (root-mean-square displacement of 0.13 pixels or 0.7 μm (Supp. Fig. 7)). These shifts were corrected using NoRMCorre^22^ before footprint extraction.

The neural activity recorded by the Kiloscope in one mouse was further analyzed as follows. Extracted neural activity was sorted by non-negative matrix factorization (NMF) factor to highlight potential low dimensional components (Fig. 3d). Note that the correlated activity visible in each component is driven by footprints from across the FOV (Supp. Fig. 10). We trained an SVM model on half of the data and used the other half to decode the mouse’s combined position and movement direction, with the resulting accuracy at least four times chance level (Fig. 3f). Increasing the number of footprints used in the decoder yielded greater decoding accuracies, as expected (Fig. 3g). Notably, footprints in different regions of the FOV yielded substantially different decoding accuracies (Fig. 3h). The same approach was also used to decode even more fine-grained position information (Supp. Fig. 8). An examination of individual neural firing rate maps yielded footprints whose activity was localized to subregions of the maze (Supp. Fig. 9).

Overall, these microscopes afford the opportunity to record populations of neurons in freely-behaving animals previously inaccessible to functional imaging, and to acquire data in larger populations of neurons than previously possible with head-mounted microscopes. In recent years, important discoveries have been made using small model organisms such as food-caching birds, tree shrews, singing mice, and bats^24-27^. We anticipate that more sophisticated miniaturized open-source tools for single neuron physiology will allow for further progress using these animals. Access to datasets with thousands of neurons in freely moving animals will advance our understanding of how populations of neurons underlie natural behaviors, such as navigation and social interaction, that are difficult to study in head-fixed preparations^9,10^.

## Supporting information

Representative video from zebra finch

Representative video from mouse

## Acknowledgements

We thank Daniel Aharoni and Jonathan Newman for many helpful discussions and for their technical expertise. In addition we thank Matthew Wilson, Jonathan Newman, Jakob Voigts, Nader Nikbakht, Andrew Bahle, Michael Happ, Alyx Tanner, and Joergen Kornfeld for comments on the manuscript. We thank Wei Guo for providing transgenic mice and Qianli Xu for baseplate designs and for running mice. M.S.F. acknowledges funding through the McKnight Foundation. J.R.S. acknowledges funding through the Harold and Ruth Newman Family Hertz Graduate Fellowship. G.F.L. acknowledges funding through the Simons Foundation (grant 542977ASPI). J.J.Z. acknowledges funding through the NIH (R21 EY028381-01).

## Data Availability

The imaging data presented in this paper is available from the corresponding author upon reasonable request.

**Supplemental Figure 1.**
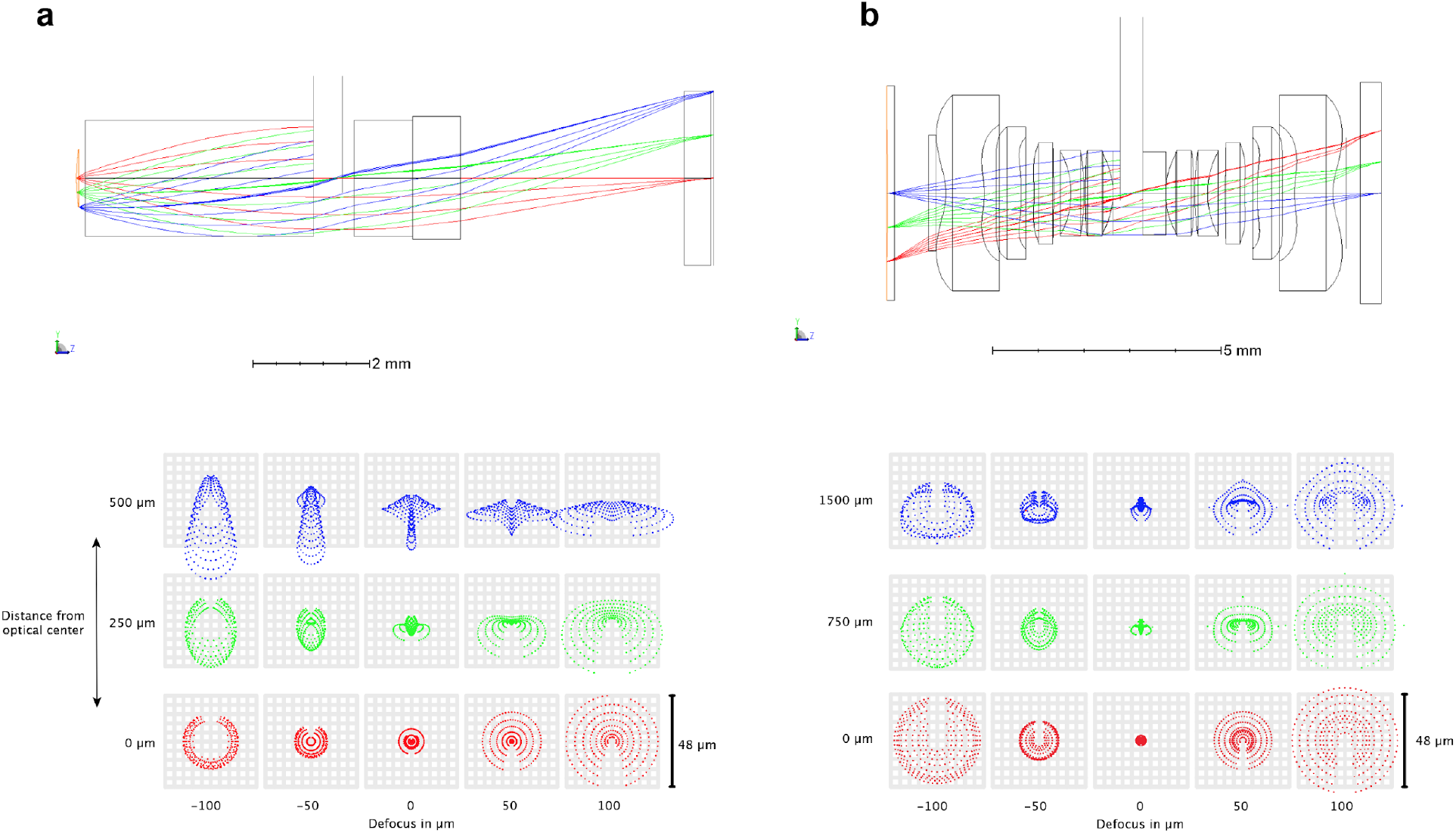
Ray trace analysis of optical assemblies. **a**, Ray trace simulation of Featherscope lens assembly (above) and point spread function calculated at the image plane (below). The grid spacing is equal to the pixel width. The cross-shaped behavior of the PSF at the edges of the field of view is characteristic of astigmatic aberrations. **b**, Same ray trace simulation for the Kiloscope lens assembly.

**Supplemental Figure 2.**
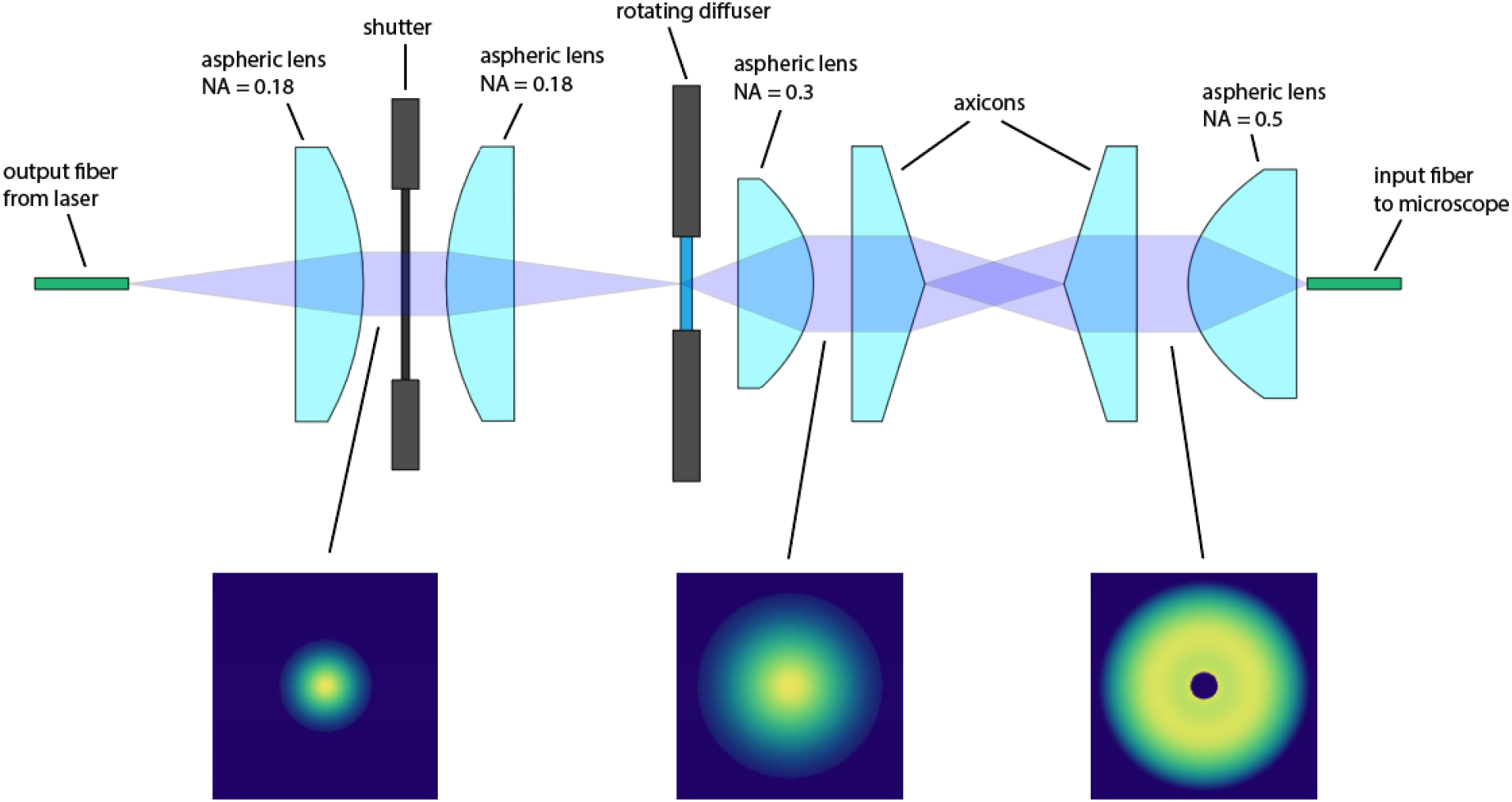
Illumination coupling optics. This optical assembly adapts the output NA of the laser to the NA of the microscope objective, scrambles the modes to prevent illumination speckle, flattens the illumination profile, and provides computer control over illumination to prevent photobleaching. First, a pair of aspheric lenses relays laser light through a computer-controlled shutter onto a rotating diffuser that adds a random phase modulation to the beam. Then, another aspheric lens collimates the output of the diffuser through a pair of axicons that radially invert the beam intensity profile, thereby flattening it. Finally, a high NA aspheric lens couples the beam into the input fiber of the microscope. The beam profile is illustrated below at various points in the assembly. Spacing between the axicons was adjusted for optimal microscope performance, and it was noted that the optimal configuration resulted in a central dark spot in the output beam. This spot is not present at the object plane due to imperfections in the fiber.

**Supplemental Figure 3.**
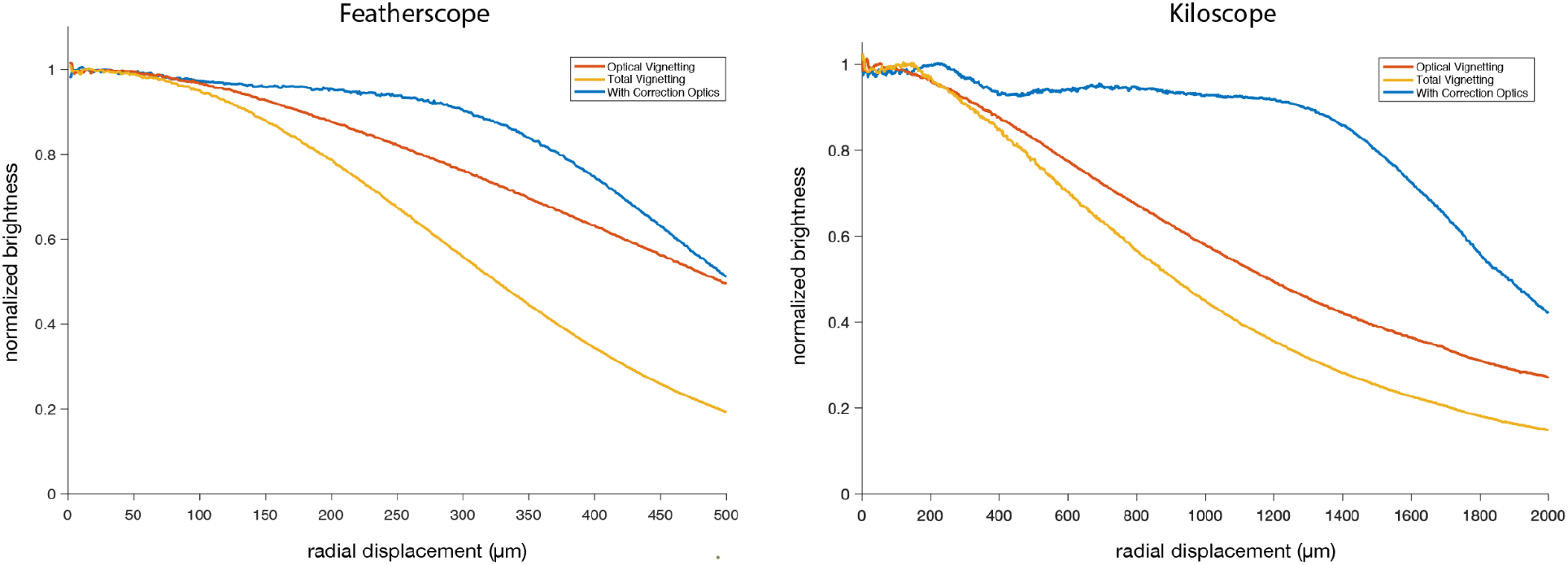
Vignetting correction. Radial profile of the signal collected while imaging a thin layer of fluorescent dye using different optical configurations. Optical vignetting (orange) illustrates the effects of light losses within the microscope optics and was measured by illuminating the dye with a diffuse light source opposite the objective. Total vignetting (yellow) illustrates the decrease in fluorescent signal caused by the combination of optical vignetting and a Gaussian-shaped excitation light profile. The curve with correction optics (blue) illustrates the flat signal intensity profile that can be achieved with an axicon pair that increases the excitation intensity at the edges of the field of view.

**Supplemental Figure 4.**
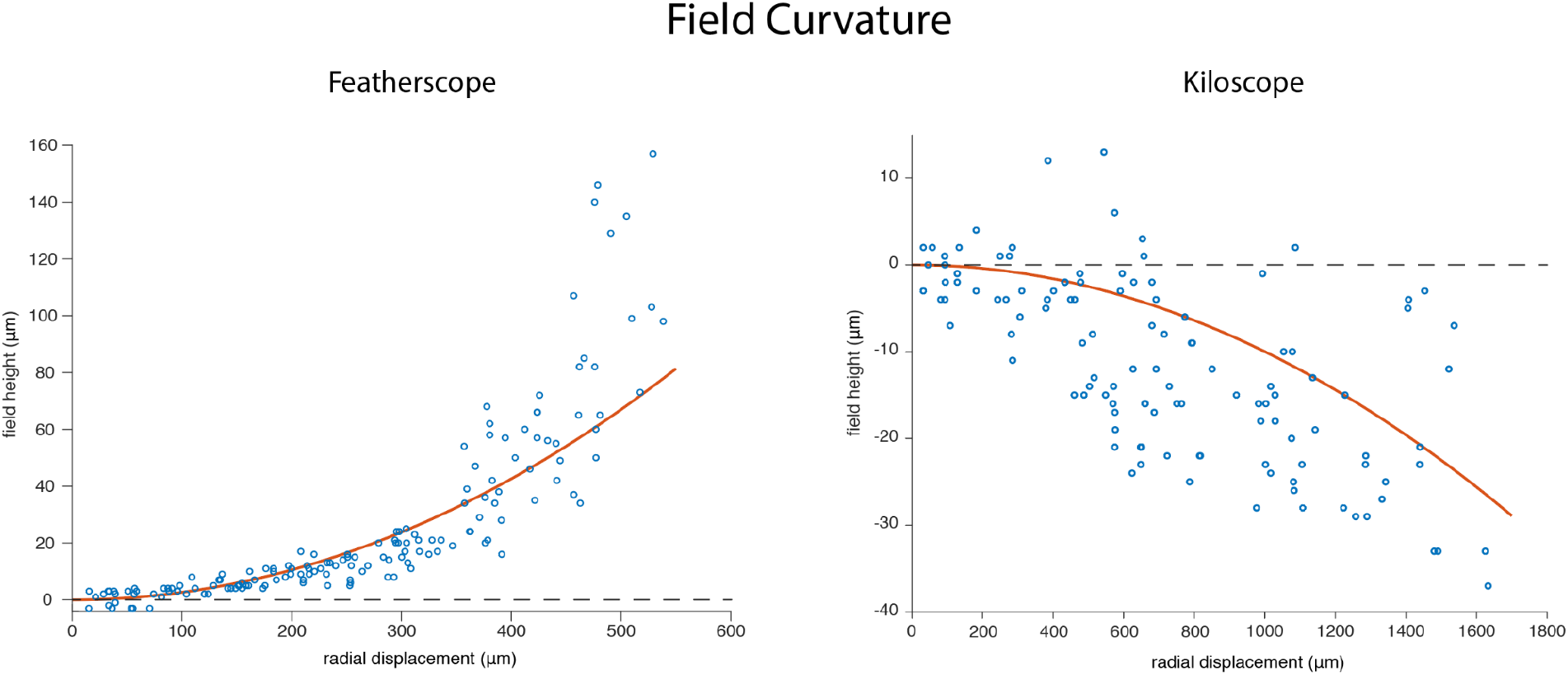
Field Curvature Measurements. Results of bead PSF measurements compared to simulations. Bead height at best focus is plotted in blue for the set of beads used in the PSF measurement (Fig 2a,d). The observed field curvature is compared to simulation results for a spherical surface of best focus, plotted in orange.

**Supplemental Figure 5.**
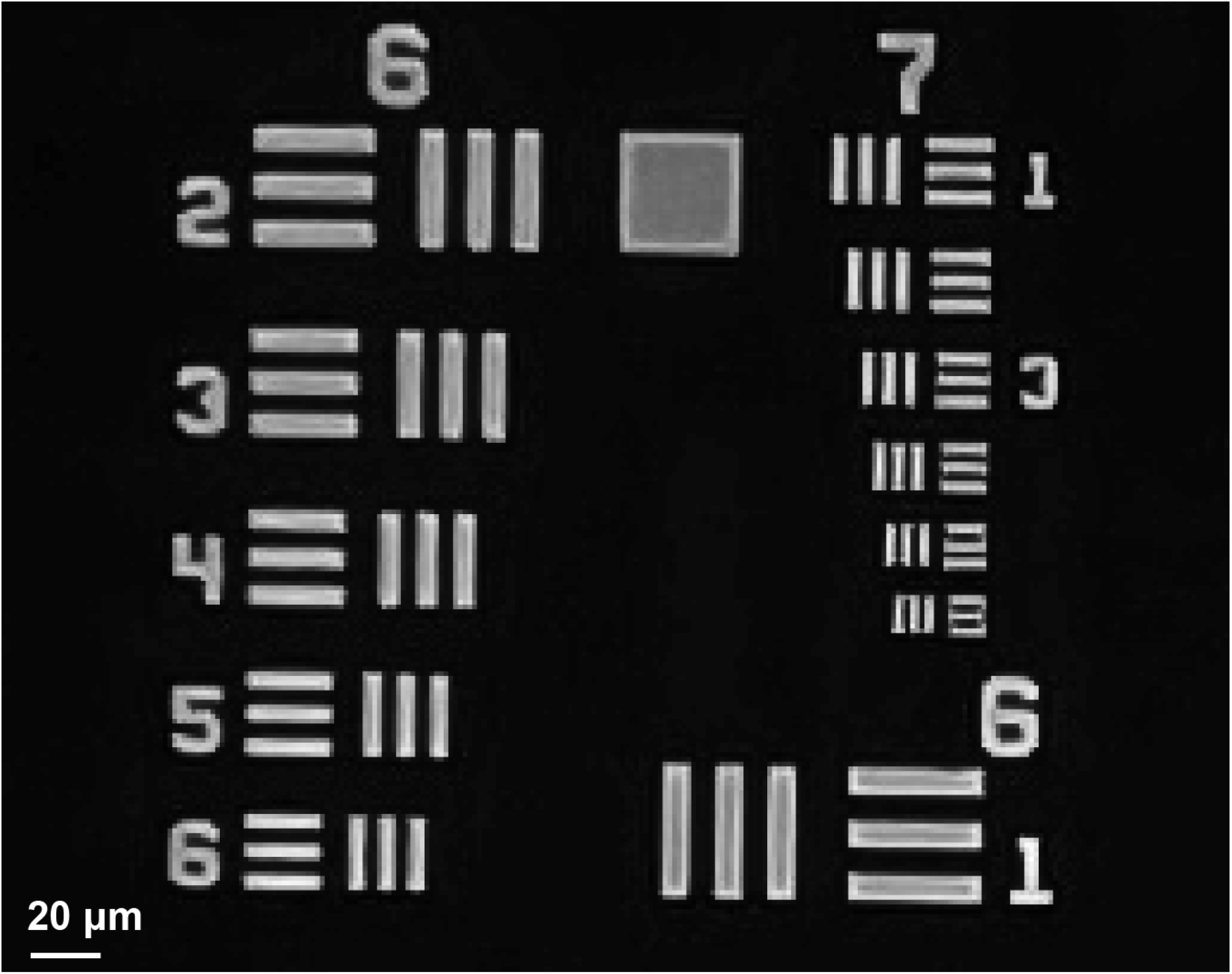
Ultimate attainable resolution of Kiloscope optics. Image of USAF resolution target using a Sony IMX260 image sensor with 1.4 μm pixel width. The smallest element in group 7 is resolvable and has a linewidth of 2.2 μm.

**Supplemental Figure 6.**
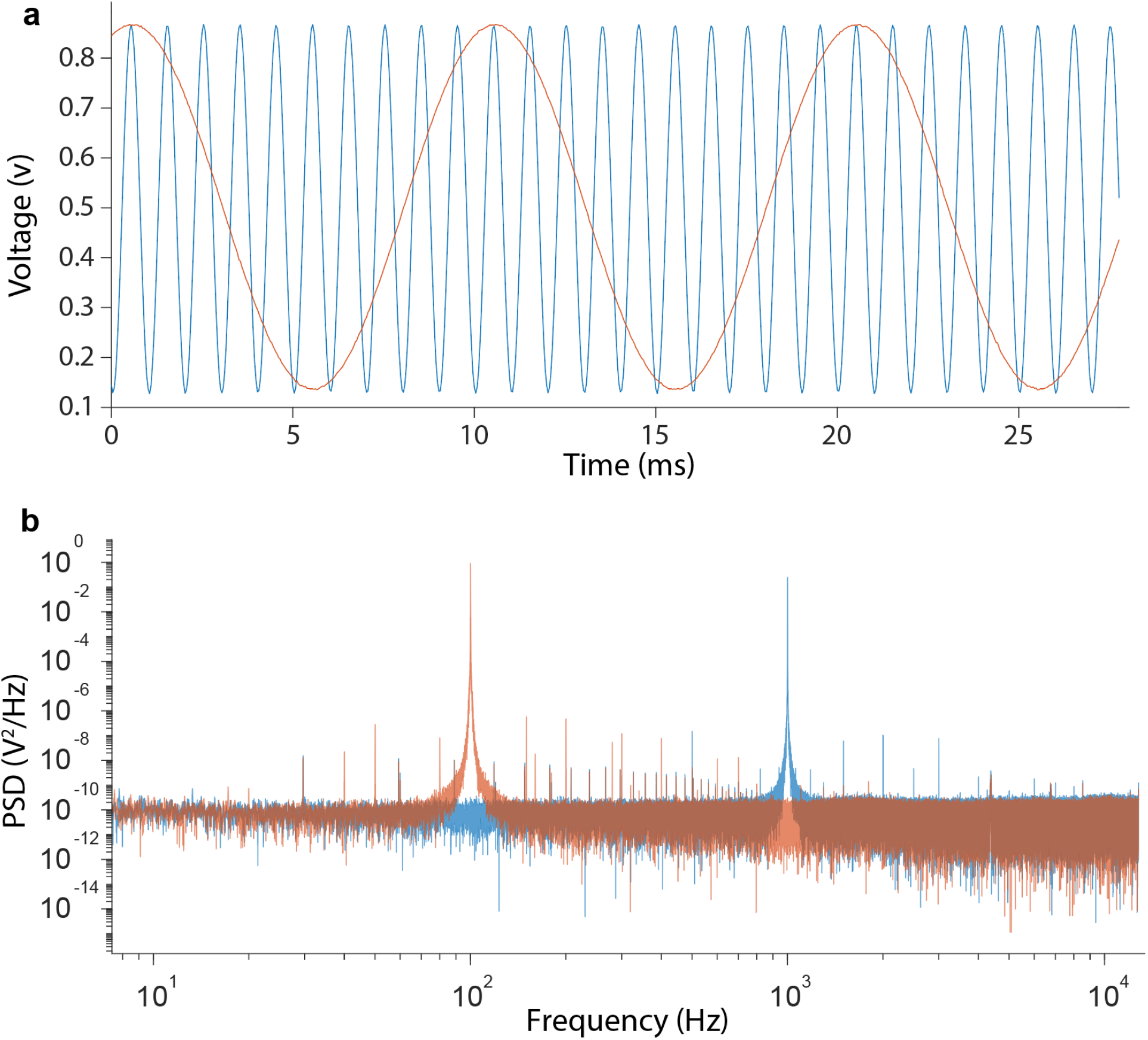
Analog-to-digital converter functionality. **a**, Time series recording of two test signals at 100 Hz (orange) and 1000 Hz (blue) using the onboard analog-to-digital converter, sampling at 25.5 kS/s for each signal. **b**, Power spectral density of the test signals. The spectrum was calculated using Thomson’s multitaper estimate with Slepian tapers. The time series deviates from a pure tone with a root-mean-square residual of 0.2% of the full scale, calculated using the full bandwidth of the recorded signal.

**Supplemental Figure 7.**
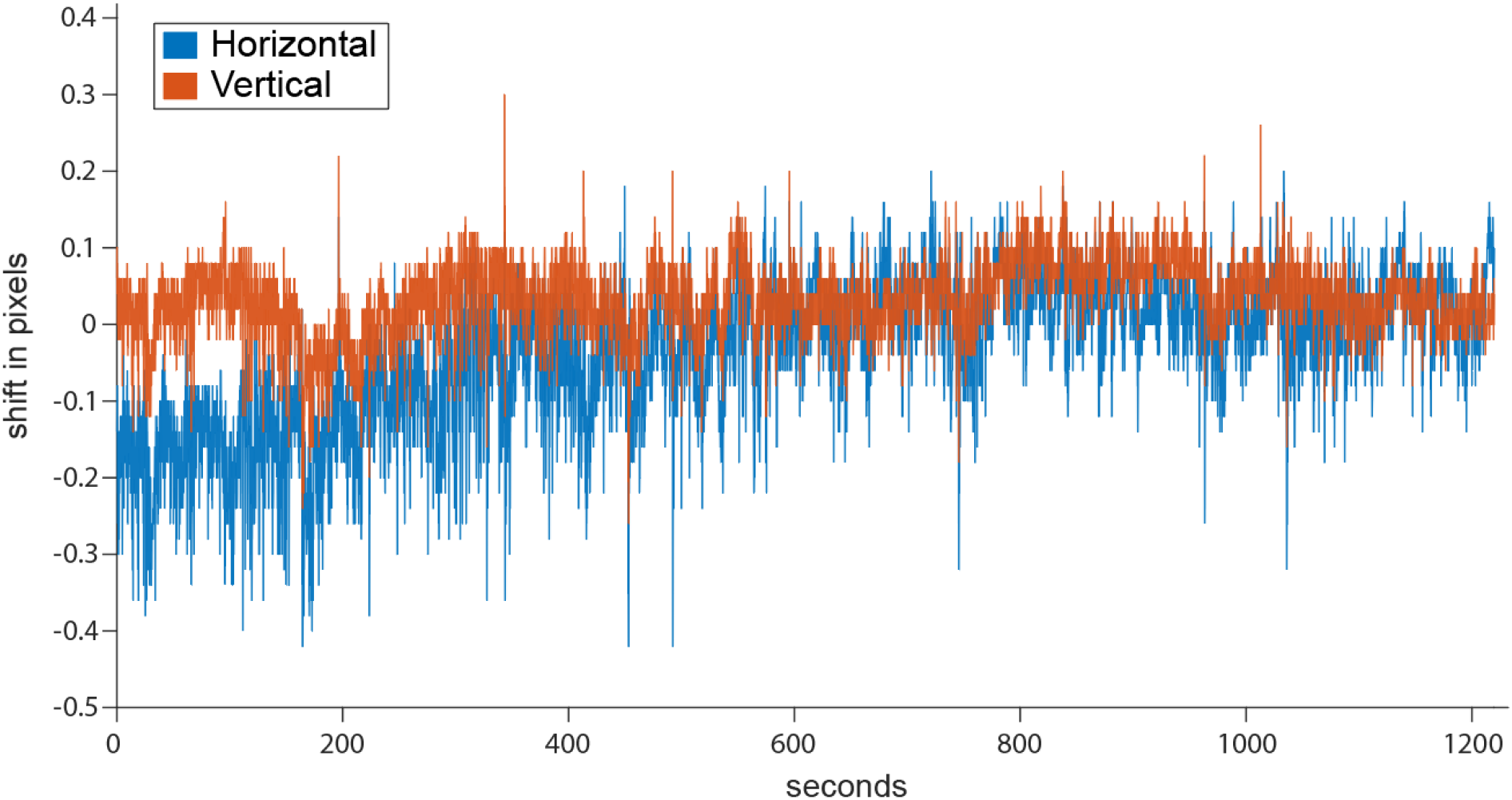
Movement observed using the Kiloscope. Time series of horizontal and vertical displacement of the FOV, calculated using NoRMCorre, during the recording shown in Fig. 3.

**Supplemental Figure 8.**
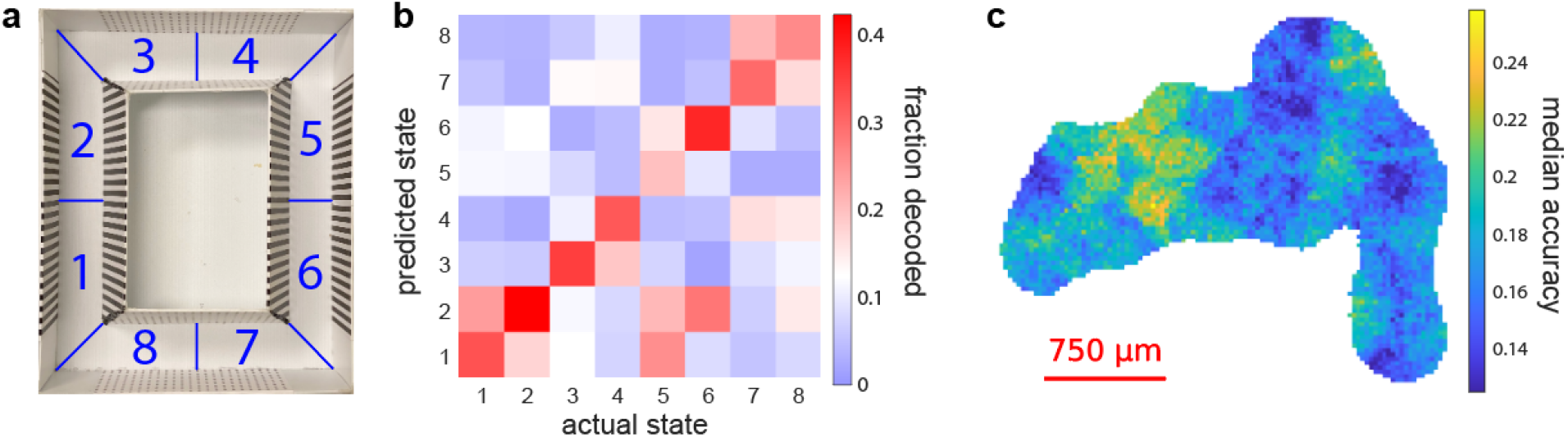
Decoding position within the maze with octant resolution. **a**, Circular maze with octants labeled; in this case, direction is not coded. **b**, Decoding matrix for a set of binary SVM models trained to predict the labels in (a) based on the activity in Figure 3d. White is set at chance level. **c**, Map of decoder accuracies using the 64 closest neurons to a given point. Colorbar minimum is set at chance level.

**Supplemental Figure 9.**
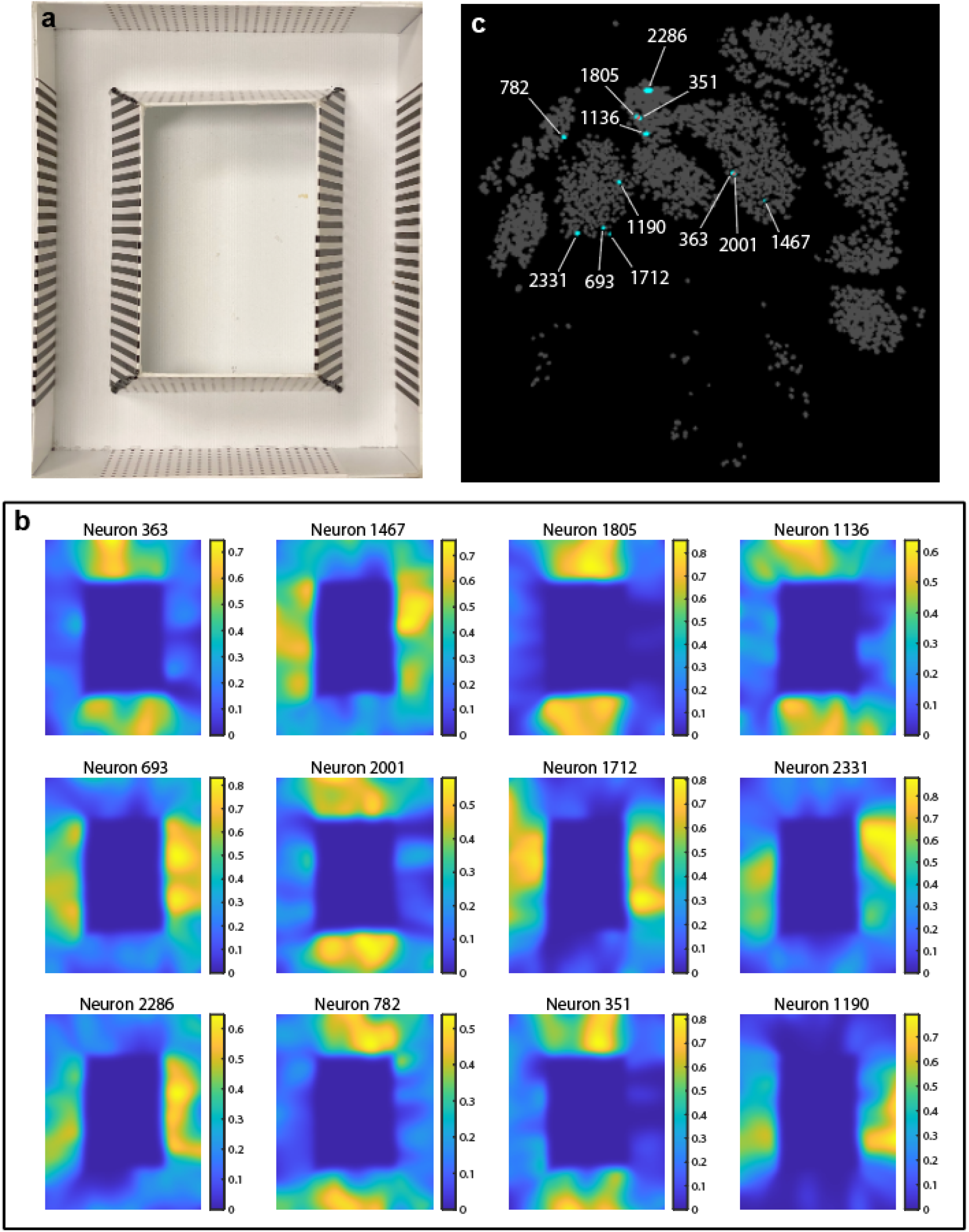
Receptive fields of selected footprints from the Kiloscope. **a**, the circular maze that the mouse explored during the recordings in Fig. 3a-d. **b**, receptive fields plotted for a collection of footprints from Fig. 3a-d. The plotted intensity is the fraction of the time a given footprint was active at a given location. Footprints were selected by eye for large-scale structure in their receptive fields. Occupancy within the track was approximately uniform, as indicated in Fig. 3c. **c**, locations of the selected footprints within the FOV. Footprints were selected blindly without knowledge of their location within the FOV, but after unblinding most footprints were found to cluster in a subregion of the FOV associated with higher spatial decoding accuracy (Fig. 3h and Supp. Fig. 8c). Regions of overlap between footprints are plotted in red.

**Supplemental Figure 10.**
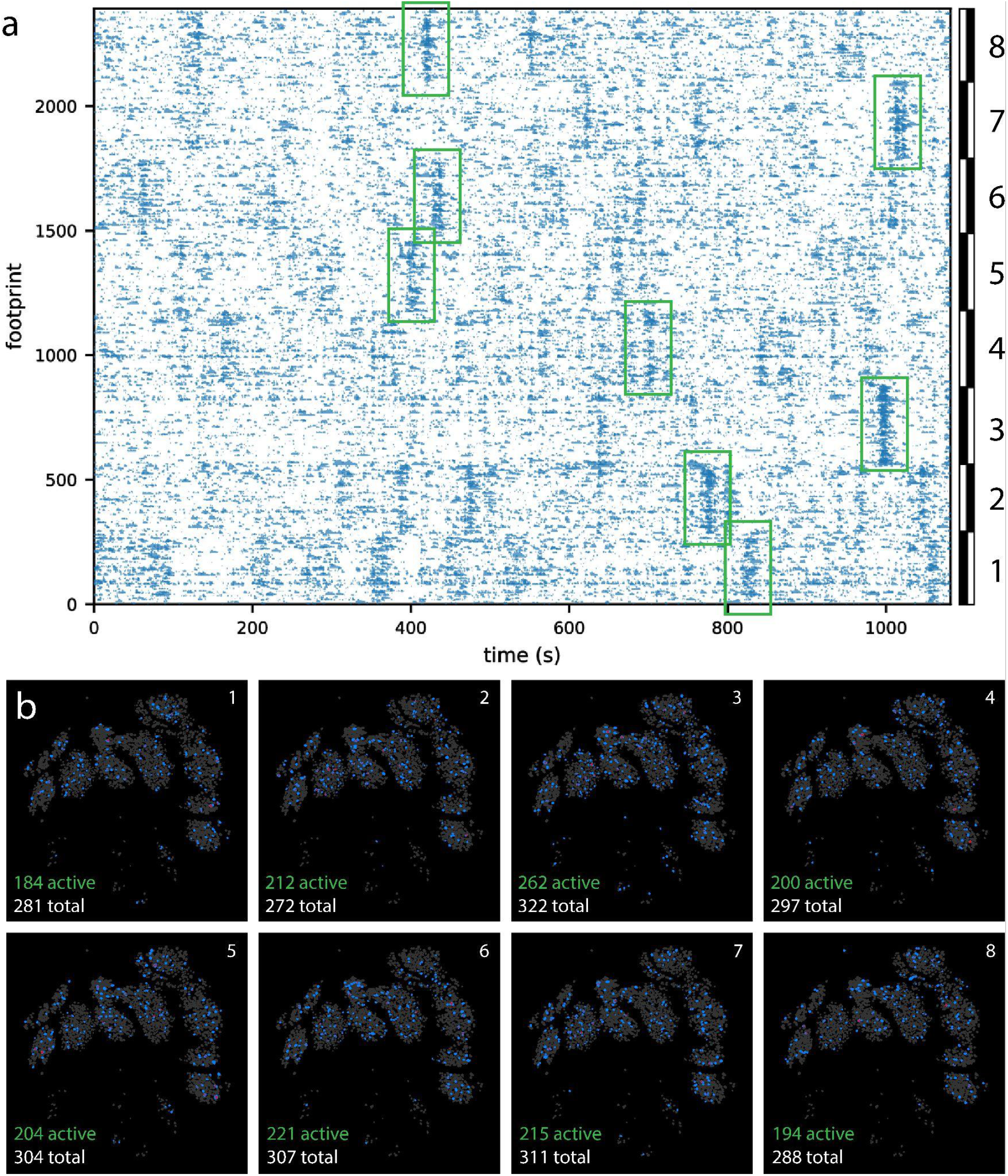
Spatial distribution of coactive neural populations in the Kiloscope dataset shown in Figure 3. **a**, Ridgeline plot from Fig. 3d; highlighted in green is a single large correlated event for each NMF component. **b**, Locations of footprints that participate in each highlighted event. Pixels containing one active footprint are colored blue, two or more overlapping footprints colored red, and inactive footprints colored gray. Footprints are thresholded at the 10% level. Listed in each panel are the number of footprints active in the associated event and the total number of footprints in the NMF component. Note that each event involves footprints distributed across the entire field of view, demonstrating that correlated events are not due to individual neurons split into multiple footprints.

**Supplemental Figure 11.**
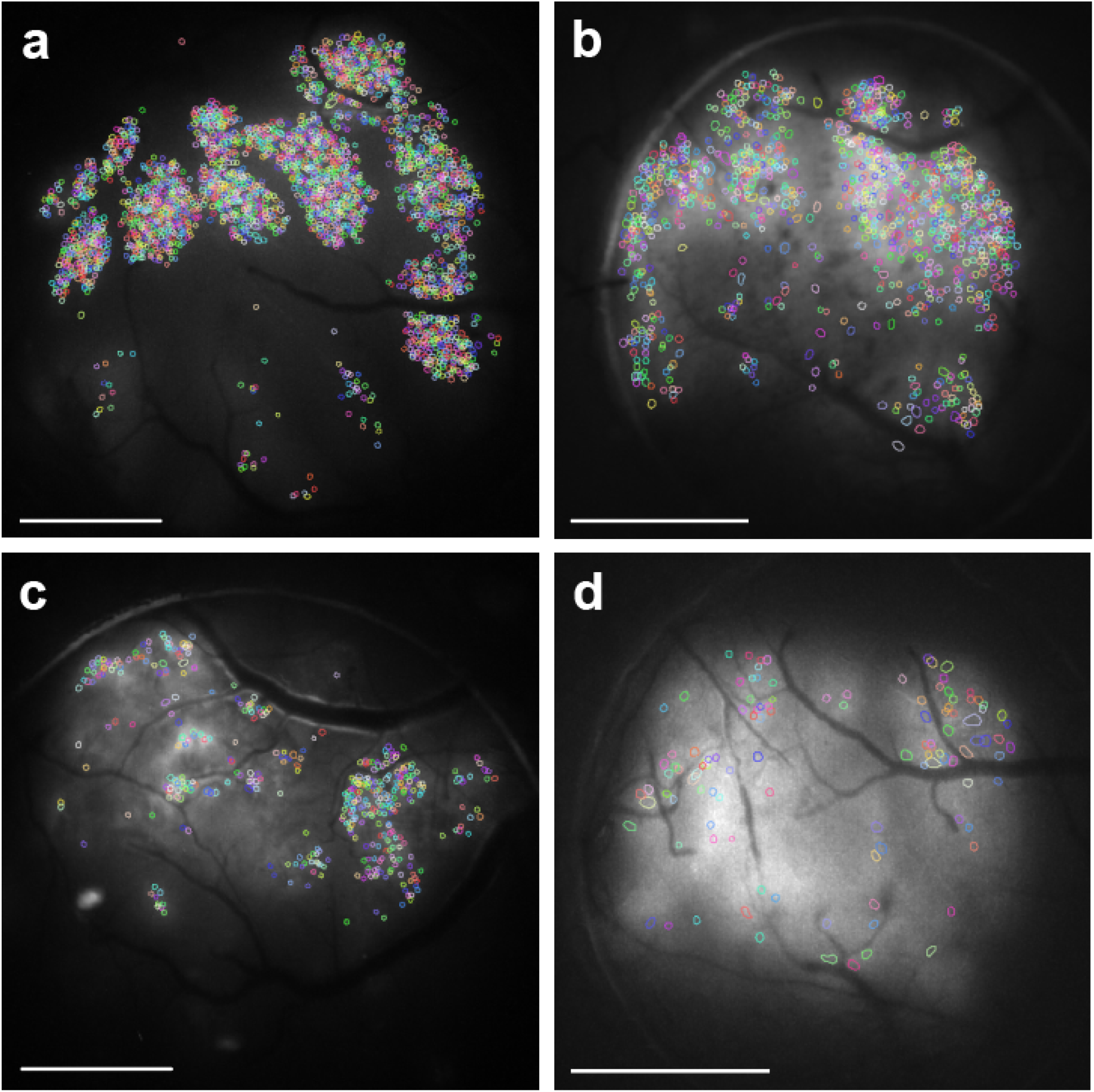
Extraction results from the four mice presented in the paper. Footprints for each extraction are plotted as colored outlines against a grayscale background of the maximum intensity of median filtered data over the course of a recording session. Footprints were extracted using EXTRACT and manually curated for panel (a) based on videos of each footprint’s activity. For panels (b-d), we inspected each footprint by eye and removed visually anomalous footprints and footprints overlapping with blood vessels. The high variability in the number of extracted footprints is likely due to variations in virus expression levels between individuals. All scale bars are 1 mm. **a**, 4 mm cranial window, 2382 footprints after curation. **b**, 3 mm cranial window, 1029 footprints. **c**, 4 mm cranial window, 401 footprints. **d**, 3 mm cranial window, 112 footprints.

**Supplemental Figure 12.**
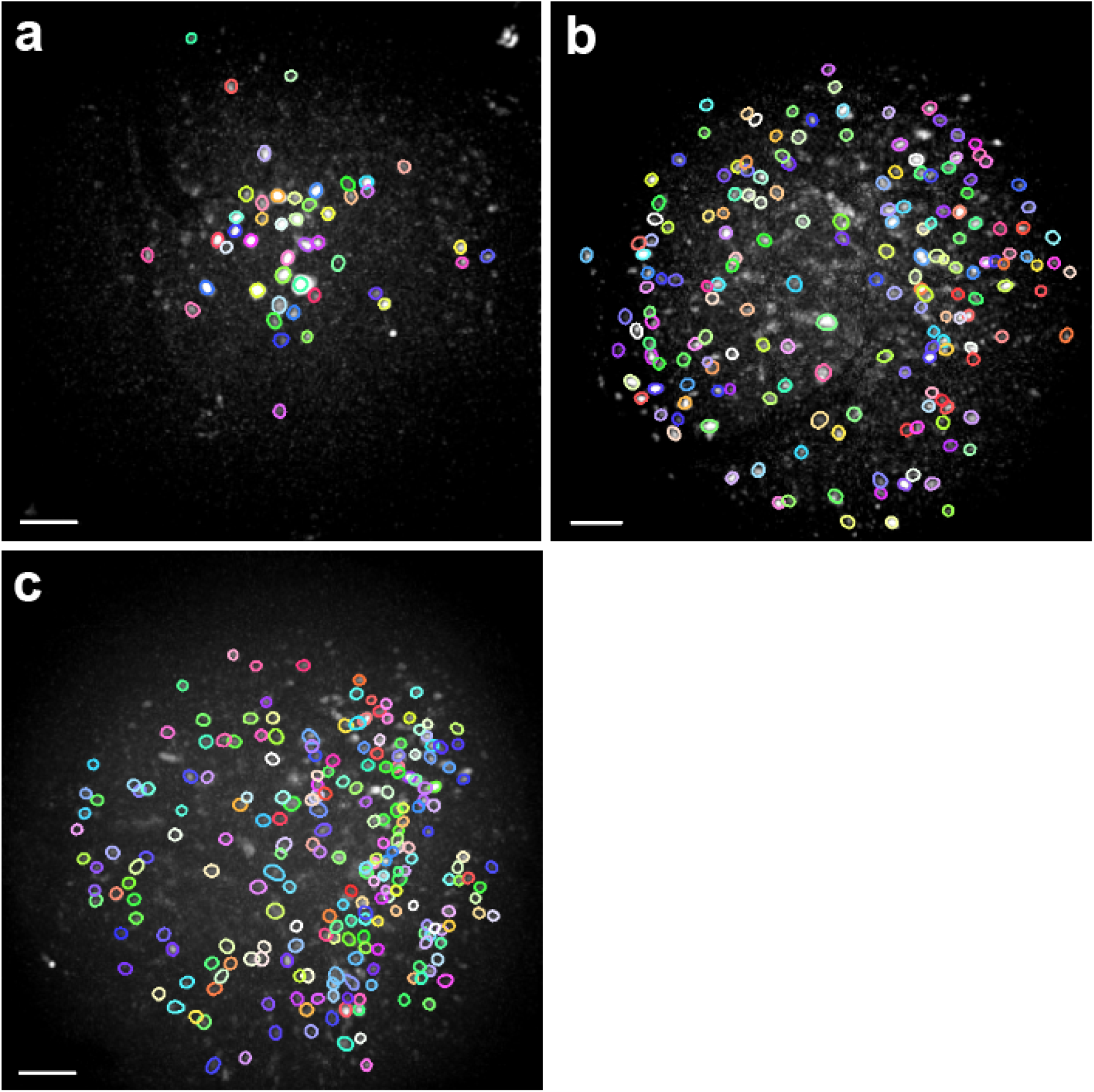
Extraction results from the three birds presented in the paper. Footprints for each extraction are plotted as colored outlines against a grayscale background of maximum intensity of median filtered data over the course of a recording session. Footprints were extracted using EXTRACT and manually curated based on videos of each footprint’s activity. The high variability in the number of extracted footprints is likely due to variations in virus expression levels between individuals. All scale bars are 100 μm. **a**, 46 footprints after curation. **b**, 177 footprints after curation. **c**, 225 footprints after curation.

## Methods

### Electronics

The imaging electronics are designed around the Python 480 image sensor (ON Semiconductor NOIP1SN0480A for Featherscope and NOIP1SP0480A for Kiloscope). Our basic data acquisition architecture is modeled after the UCLA Miniscope system^1^. The image sensor outputs data on a 10 bit parallel bus that is sent over coax to a data acquisition (DAQ) board using a serializer and deserializer pair (SERDES; Texas Instruments DS90UB913A-Q1 and DS90UB913A-Q1). The DAQ board uses a Cypress CYUSB3014 controller to send the image data over a USB 3.0 bus to an acquisition computer. The DAQ board is programmed to present itself as a USB video class device (UVC) with uncompressed RGB565 video frames, as 10 bit monochrome frame formats are not supported by UVC. The 10 bit monochrome pixel data occupies the lower 10 bits of the RGB565 pixels (blue and red bits), and conversion from RGB565 to 10 bit monochrome pixel formats is done offline using custom software.

Our image sensor PCB also includes a microcontroller (Atmel ATtiny816) that is used to configure the Python 480, perform power sequencing, provide two channels of analog to digital conversion (ADC), and act as a bridge between the I2C bus provided by the serializer and the SPI bus on the image sensor. We use the microcontroller’s onboard ADC to read two analog signals. The digitized reads are transmitted as USART serial data over the two most significant bits of the serializer’s 12 bit data bus. On the DAQ side, we split the USART bus from the output of the deserializer, convert it to two low-voltage differential signal (LVDS) pairs, and send them off-board via an HDMI connector. We can then acquire these ADC signals with the acquisition computer using an Arduino “Teensy” microcontroller.

The image sensor PCB was designed to be as compact and lightweight as possible. It is a six-layer PCB weighing approximately 300 milligrams. Because it is so small, heat management has a significant effect on image sensor operating temperature and thermal noise. Following the NINScope system, we have disabled the image sensor PLL to reduce power consumption^2^, and have attempted to maximize PCB thermal conductivity by coupling the image sensor and serializer to the ground plane using dense vias. An optional copper fin heat sink can be soldered to the PCB that weighs 150 milligrams and improves SNR by a factor of approximately three.The Kiloscope data was taken with such a heatsink installed, but the Featherscope data was taken without any heatsink.

The DAQ board is a modified version of the UCLA Miniscope v3 DAQ board that uses all twelve bits of the SERDES parallel data bus, rather than the eight in the original design. Two of the four additional bits are routed to the Cypress and used to acquire the full 10 bit output of the image sensor. The other two bits encode ADC reads and are routed to an LVDS converter. From there, the differential signals are sent to an off-board Arduino that sends them over USB to the acquisition computer.

### Optics and Construction

#### Featherscope

The Featherscope is built from microoptical components cemented into a 3D printed body (Rosenberg Industries LLC) using UV-cure optical cement (Norland Optical Adhesive 61 and 63). The body consists of two subassemblies that are attached once the optics are installed. Body components are printed using Formlabs Black resin, known to exhibit low autofluorescence^3^.

The lower subassembly holds the GRIN objective lens (Edmund Optics #64-537), coupling prism (Edmund Optics #66-770), and custom fiber bundle (Acrolite, Inc.). Once the objective lens is installed in the body, the prism and fiber bundle are cemented to the back face and covered in a drop of black nail polish (OPI Black Onyx). The nail polish is allowed to dry, and any excess is then carefully trimmed away with a scalpel to expose as much of the back aperture of the objective as possible while maintaining enough thickness (∼50 μm) to block excitation light leakage. A machined magnetic steel flange is cemented to the bottom face of the body to allow for mating with commercially available magnetic baseplates weighing 230 mg (Inscopix).

The upper subassembly holds the tube lens, optical filters, focus mechanism, and image sensor PCB. The tube lens is a GRIN lens (Edmund Optics #64-537) ground down to a total length of 1.00 mm (Pioneer Precision Optics). The tube lens, dielectric filter (3 × 3 × 0.5 mm, Chroma ET525/50m) and Wratten filter (No. 8, Knight Optical #494FWP) are attached to the body using optical cement. A foam spring is then inserted. The spring is made from compressible polyurethane foam (McMaster #86375K121) with a 3.0 mm hole punched using a biopsy punch. The image sensor board is then installed and held in place with a focus bracket. The bracket is constructed from 0.004” thick 316 stainless steel with a #000 nut cemented to the underside using JB-Weld epoxy. A hole is punched above the nut and a #000-120 screw is threaded into the bracket. During normal operation, the screw pushes against the top of the onboard microcontroller and compresses the foam spring.

Once the two subassemblies are built, they are attached together using optical cement. The fiber bundle is then attached to the relief arm of the upper subassembly body using dental floss and potted using 5 minute epoxy (Loctite 1365868). Thin and flexible lines for power (Cooner Wire #CZ1174) and coaxial data (Cooner Wire #CW6563) are braided together with the fiber bundle and secured using dental floss.

#### Kiloscope

the Kiloscope is constructed in a similar manner to the Featherscope using two subassemblies of optics mounted in 3D printed housings. The lower subassembly holds the objective lens, an imaging lens extracted from the rear camera module of a Samsung Galaxy S9 smartphone. These imaging lenses are carefully removed from rear camera replacement modules available from third-party vendors. The lens is installed in the printed housing and enclosed at the top and bottom by two pieces of protective No. 1 coverglass. The fiber-prism assembly is installed at the top coverglass using the same procedure as the Featherscope.

The upper subassembly holds the dielectric filter, tube lens, focus mechanism, and imaging PCB. The tube lens is identical to the objective lens and is attached directly to the dielectric filter. Compressible sorbothane rubber strips (50OO Durometer, McMaster #8824T11) are used as focus springs and are inserted on ledges designed into the housing. The PCB is then installed and held in place with a focus bracket similar to that used with the Featherscope. The fiber is secured to the relief arm of the housing and is braided together with the electrical lines. The microscope slides into a custom machined aluminum baseplate weighing 300 mg.

The Kiloscope exhibits some degree of autofluorescence caused by the plastic optics in the objective lens assembly. However, this fluorescence bleaches out to a negligible background level after an initial exposure period.

Weight measurements for both microscopes were made by weighing each microscope with the tether attached and supported horizontally at the same height as the microscope. This method was used because the tether is permanently attached to the microscope during construction. This measurement technique thus includes a small portion (∼5 cm) of the tether weight as well.

### Imaging Performance Characterization

Imaging performance was measured using two methods. The first was using a standard 1951 USAF resolution test target against a fluorescent background. This method has the benefit of intuitive interpretation and broad recognizability, but it is somewhat subjective and does not include information about the resolution of the system at different points in the FOV. Miniature endoscopes such as ours have significant aberrations that vary across the field of view and tend to be minimized at the center, so measurements using resolution test targets at a single position can overestimate the full-field resolution.

Because of these limitations, we decided to quantify the point spread function of our microscopes using an array of beads imaged at various positions and vertical displacements within the FOV. The PSF contains the same information as a two-dimensional modulation transfer function (MTF) measurement taken at various positions, but has the added benefit of being intuitively interpretable for researchers who are not familiar with optical engineering methods.

For the PSF data in Fig. 2, we imaged a slide of 1 μm diameter fluorescent beads at a series of vertical positions spanning 100 μm of depth. We then found the position corresponding to the most focused image for each bead, and fit a two dimensional Gaussian distribution to that image to find an approximate FWHM. We also measured the field curvature by plotting the vertical position of best focus as a function of radial distance (Supp. Fig. 3). We sampled 155 and 118 beads for the Featherscope and Kiloscope analysis, respectively.

### Commutator

Our commutator simultaneously provides power, RF data, and illumination light to our microscopes. We couple DC power and RF data through an electrical slip ring (Moflon MSP1069) similar to existing open source designs (University of Colorado ONE Core^4^). We transmit light through an optical fiber threaded through the bore of the slip ring and attached to an optical rotary joint (Thorlabs RJ1) mounted coaxially with the slip ring. We drive the commutator using a motor controlled by a magnetic feedback signal, similar to existing designs^5^.

The slip ring presents an impedance mismatch for the 50 Ohm transmission line and on its own causes disruptive signal reflections. However, because the slip ring size is a small fraction of the carrier wavelength, we can partially compensate for the mismatch by adding a small tuning capacitor (2-6 pF) in parallel with the slip ring. Using this capacitor, along with boosting the forward channel low frequency gain of the SERDES, we can transmit data through the slip ring without excessive frame dropping.

### Coupling Optics

The illumination coupling optics are based on a set of four aspheric lenses (Thorlabs C280TMD-A, C230TMD-A, and A397-A) that adapt the laser output NA of 0.1 to the higher microscope fiber NA of 0.5. A shutter placed in between the first two lenses provides fast control over illumination to prevent photobleaching while avoiding power instability caused by modulating the laser current.

We reduce speckle in the laser output using an oscillating diffuser (Optotune LSR-3005-17S-VIS) placed at the imaged laser spot between the second and third lenses. This diffuser adds a random time-varying phase to the beam waist that randomizes the speckle pattern in the object plane. The diffuser oscillates at 300 Hz, so a single frame integrates over many of these random patterns, averaging them together and reducing the overall speckle. We do not observe any residual speckle in our *in vivo* data after despeckling.

The Gaussian output of the laser results in a Gaussian distribution of illumination in the object plane. This distribution combined with the optical vignetting of the microscope results in dim fluorescence signals and thus low SNR at the edges of the FOV. To compensate, we use a pair of axicons (Thorlabs AX1220) between the third and fourth lenses that transforms the Gaussian input into a relatively flat output. The precise shape of the output distribution can be adjusted by varying the distance between the axicons.

An illumination power of 200-300 μW was used to record data using the Kiloscope, which results in an average sample irradiance of 16-24 W/m^2^. This illumination is comparable to that of published microscopes that use 12-55 W/m^2^ to image GCaMP6f in mice^2^. An illumination power of 2.0 mW was used to record in birds using the Featherscope, which corresponds to an average sample irradiance of 790 W/m^2^. This value is significantly higher than the illumination needed to image in mice, but is equivalent to the power we have used with Inscopix systems in adult birds. We believe this discrepancy results from the lower level of viral expression that we can achieve in birds with viral vectors typically used in mammals.

## Data Analysis

In vivo imaging data was analyzed using the NoRMCorre and EXTRACT algorithms^6,7^. For the Kiloscope data, we first ran NoRMCorre to correct for brain motion. We then ran EXTRACT using mostly default parameter values but with an increased value for the minimum cellfinding SNR of 2. For the mouse with the best calcium indicator expression, this extraction yielded 2948 footprints, which were then manually curated by viewing the median-subtracted video in the region of each extracted footprint at times corresponding to the peak extracted activity. We scored each footprint by whether it appeared by eye to be a spatially distinct footprint modulated in time and whether it was spatially separated from blood vessels. This manual curation yielded 2382 putative neurons and is a lower bound on the actual number of neurons in the dataset.

The resulting time series were factored using nonnegative matrix factorization (NMF) into eight component factors, and the data plotted in Figure 3d was sorted according to these factors to highlight potential low dimensionality of the data. Activities very close to zero are not plotted for visual clarity.

For the Featherscope data, we extracted footprints using the same pipeline, with the exception that image alignment was performed using custom software. The extracted footprints were manually curated in the same way; for the bird with the best calcium indicator expression, this curation yielded 177 putative neurons used in the analysis. These footprints are plotted in Figure 3l, with activities very close to zero not plotted for visual clarity.

Song production during Featherscope experiments was quantified by counting the number of syllables each bird produced each day. On many days birds sang very little because they were handled for focus adjustment or hardware modifications, so the rates cited in the paper reflect the maximum singing rate per day for each bird.

The decoding analysis in Figure 3e-h and Supp. Fig. 8 was carried out using a set of binary SVM classifiers in a one-versus-all encoding scheme. The model was trained using a random contiguous 50% of the data with temporal circular symmetry and tested on the other 50%. We omitted 1% of both the training and test data, at the transition between training and test data, to avoid overfitting the model on the data adjacent to the boundary. Decoding matrices were calculated by averaging the performance of many decoders on different random holdouts.

For each decoder size bin in Figure 3g, a hundred different random subsets of footprints were used to generate corresponding decoding matrices for each subset. The median decoding accuracy reported in Figure 3g is the median of the diagonal elements of each decoding matrix. For the maps in Figure 3h and Supp. Fig. 8c, decoders were trained using the 64 footprints closest to each pixel. Pixels were omitted from the analysis if there were fewer than 64 footprints within a 270 μm radius of the pixel.

### Video Processing

Example videos were generated using custom Julia software. The fluorescence signal was median filtered over time. For the mouse data (Supp. Video 2), the signal was temporally downsampled by a factor of four. The image levels of both videos were adjusted for display purposes.

Maximum intensity projections (Fig. 3a, e) were calculated using custom Julia software. The fluorescence signal was first median filtered over time and then spatially bandpass filtered. The image levels of each maximum intensity projection were adjusted for display purposes.

### Subjects and Calcium Imaging

Calcium imaging was performed in juvenile male zebra finches (*Taeniopygia guttata*) 50-70 days post-hatch (dph) during undirected song, and in adult C57/B6 mice during free exploration of a circular maze. Birds and mice were obtained from the Massachusetts Institute of Technology breeding facility (Cambridge, Massachusetts) and Jackson labs, respectively. Video and behavioral data was collected using custom Bonsai workflows^8^. The care and experimental manipulation of the animals were carried out in accordance with guidelines of the National Institutes of Health and were reviewed and approved by the Massachusetts Institute of Technology Committee on Animal Care.

Prior to surgery, birds were isolated from their tutor at 35 dph. Birds were anesthetized with 1-2% isoflurane in oxygen and placed in a stereotaxic apparatus. LMAN was mapped using antidromic identification from RA as previously described^9^, and RA was injected with cholera toxin subunit B (recombinant) Alexa Fluor 647 conjugate (Invitrogen) to retrogradely label LMAN for histological verification. LMAN was then injected with AAV9.CAG.GCaMP6f.WPRE.SV40^10^ (University of Pennsylvania Viral Core) using a Nanoject II (Drummond). After a period of time to allow the virus to perfuse the neural tissue, we lowered a GRIN prism relay lens (Inscopix) anterior to LMAN, and attached it to the skull using previously described methods^11^. 10 days following the injection procedure, birds were anesthetized and a base plate (Inscopix) was attached to the skull.

Mice were anesthetized with 1-2% isoflurane in oxygen and placed in a stereotaxic apparatus. V1 was localized using stereotaxic coordinates, and a 4 mm diameter craniotomy was made to expose V1 as well as brain regions anterior and lateral to it. We then performed a series of injections of AAV1 pGP-AAV-syn-jGCaMP7f-WPRE^12^ (Addgene) with 500-600 μm spacing across the craniotomy. We injected 200 nL of the virus per site at a depth of 200 μm below the dura. A cranial window composed of one 4 mm diameter coverslip glued to a 5 mm diameter cover slip (Carolina) was inserted into the craniotomy and attached to the skull. After about two weeks to allow for expression, a custom base plate was attached to the skull.

